# Oligodendrocyte progenitor cell responses to inflammatory demyelination with aging in mouse model of multiple sclerosis

**DOI:** 10.1101/2025.09.22.677854

**Authors:** Emily E. Fresenko, Camilla N. Bahri, Benjamin J. Burson, Noor F. Ahmed, Davin Packer, Benjamin J. Tabor, Bogdan Beirowski, Wenjing Sun, Michelle A. Wedemeyer, Cole A. Harrington

## Abstract

Oligodendrocyte progenitor cells (OPCs) have the capacity to self-renew, differentiate, and remyelinate the CNS. Aging is associated with a reduction in the functional capacity of OPCs even in the absence of an autoimmune insult. To determine how aging affects the response of oligodendroglia to a strong inflammatory insult comparable to an immune-mediated demyelinating event in multiple sclerosis (MS), we performed adoptive transfer of young myelin-reactive Th17 T cells into young and aged OPC lineage tracing mice. After adoptive transfer, OPCs were enriched within spinal cord lesions of both young and aged mice. However differentiated oligodendrocytes (OLs) were significantly reduced after adoptive transfer. Both young and aged OPCs differentiated into mature OLs during adoptive transfer. Transmission electron microscopy revealed thinly myelinated axons without degenerative features that likely represent remyelinated axons in lesions of both age groups. Young and aged OPCs rise to the challenge after a strong auto-immune attack, suggesting that compensatory strategies permit both young and aged oligodendroglia to survive despite an inflammatory environment. Identifying pathways that promote resilience of oligodendroglia in the face of an inflammatory challenge will facilitate the development of remyelinating therapies for people with MS.

## Introduction

In the postnatal CNS, oligodendrocyte progenitor cells (OPCs) are lineage restricted progenitors with the capacity to proliferate, migrate, and differentiate into mature myelinating oligodendrocytes during development and adulthood (1–3) and in response to demyelinating injury (4,5). Multiple sclerosis (MS) is a chronic demyelinating disorder of the CNS in which an autoimmune attack of oligodendroglia and their myelin processes results in demyelinating lesions (6–8). OPCs and maturing oligodendrocytes have been identified in MS lesions from patients of all ages (9,10) suggesting that OPCs may retain the capacity to differentiate and remyelinate after a demyelinating attack despite an inhibitory inflammatory environment (8,11).

Aging is a risk factor for progressive accumulation of neurological disability in MS (12,13). Cellular aging in MS leads to senescence of immune cells and other CNS resident cell populations (13) which may influence the ability of aged OPC to differentiate and remyelinate. Physiological aging of OPCs, in the absence of an immune disorder or model, is associated with phenotypic changes in OPC ion channel expression (14), reduced proliferation and differentiation (14,15), increased cellular senescence and upregulation of immune response pathways (14–16). Developmental inhibitors of OPC differentiation, e.g. Wnt and hypoxia inducible factor-1 alpha (HIF-1a) pathways, are upregulated in aged OPCs and pharmacological inhibition of these pathways promotes differentiation of aged OPCs *in vitro* (16). In toxin-mediated demyelination models, OPC proliferation, recruitment into lesions, differentiation and remyelination are all slower in aged animals (17–19). Metformin plus alternative day fasting (15) and young macrophages (20) can partially rescue the slower remyelination kinetics in aged animals. These studies suggest that aged OPC function can be augmented to levels similar to young OPCs, and targeting mechanisms that promote aged OPC function may be a viable therapeutic strategy to promote remyelination in MS.

Formal investigations of the impact of aging on OPC performance after an autoimmune-mediated inflammatory insult have not been conducted to date. While toxin-mediated models benefit from stereotyped OPC responses, the lack of an adaptive immune response in these models may not accurately reflect the microenvironment of human MS lesions. Inflammatory models offer the benefit of a T cell mediated auto-immune process targeting oligodendroglia and myelin, better simulating an acute MS demyelinating event (6, 21). Although inflammatory models reproduce the histological features of severe disease, including early axonal degeneration, high inflammatory infiltrate, rapid severe disease course and lack of disease remission (22–24), the rapid progression and severity of clinical disease prevent the use of this model for chronic timepoints. Despite these limitations, the study of murine models with a strong autoimmune inflammatory response may uncover mechanisms that oligodendroglia utilize to adapt to a harsh inflammatory environment. Studies in both murine models (25–31) and human MS tissue (29,32) indicate the oligodendroglia adopt distinct transcriptional states in response to inflammation. These “disease-associated” oligodendroglia upregulate immune signaling pathways and may directly interact with T cells by expression of MHC class I and II molecules (30, 33–34). The function of disease-associated oligodendroglia is unclear and may be context and model dependent. Investigation of OPC properties in a strong inflammatory model will offer insight into how young and aged OPCs respond to an auto-immune demyelinating insult.

We performed lineage tracing of OPCs in the Th17 adoptive transfer model to investigate how aging affects the response of OPCs to a robust autoimmune demyelinating event. A tamoxifen-inducible OPC Cre line (*Pdgfra-Cre^ERT^*) (35) was crossed to an inducible GFP reporter line (*RCE- Rosa26^flstopflEGFP^*) (36), tamoxifen was administered to lineage trace OPCs starting two weeks prior to adoptive transfer of young myelin-reactive Th17 T cells into aged (10-14 months) and young (3-5 months) *Pdgfra-Cre^ERT^*;*RCE* recipients. The Th17 adoptive transfer model offers several advantages over other immune-mediated models including 1) control of adoptive transfer T cell age to prevent potential confounding from aged donor T cells (37) and 2) selective determination of oligodendroglial responses to a Th17 T cell mediated disease as Th17 T cells play a key role in MS pathogenesis (22, 38–39).

Following myelin-reactive Th17 adoptive transfer into *Pdgfra-Cre^ERT^;RCE* mice, we found that OPCs from both young and aged mice were enriched in spinal cord lesions. Higher OPC densities were seen in lesions compared to non-lesion white matter suggesting that the lesion environment promotes OPC recruitment and/or survival mechanisms. Differentiated oligodendrocytes (OLs) were reduced in adoptive transfer in both young and aged animals. A small minority of lineage traced OPCs differentiated into mature OLs in both ages. Transmission electron microscopy (TEM) analysis of lesions identified similar proportions of thinly myelinated axons without degenerative features in both young and aged lesions suggesting that remyelination can occur even in the presence of an ongoing inflammatory response. These findings indicate that both young and aged OPCs respond similarly to an auto-immune inflammatory demyelinating insult and OPCs may have compensatory strategies to support survival and physiological function in an inflammatory environment.

## Results

### Aged Th17 adoptive transfer animals have increased disease severity

To identify differences in the response of young and aged OPCs to an adaptive auto-immune response, young and aged mice cohorts were used for this study (Figure 1A, age at sacrifice/tissue analysis-young: P87-P148, 3-5 months; aged: P295-P427,10-14 months). Aged mice age range was selected based on mouse aging functional studies correlating our aged animal cohort to human age in 4^th^ decade (40) which is the average onset of late-onset relapsing remitting MS (LORRMS) and progressive MS (41,42). Experiments with aged animals beyond 14 months was not performed due to increased disease severity and early morbidity in our aged cohort (Figure 1B-C). OPC lineage tracing was initiated by administering tamoxifen to young and aged *Pdgfra-Cre^ERT^*; *RCE* (*Rosa26^flstopflEGFP^*) mice two weeks prior to adoptive transfer (Figure 1A) to avoid tamoxifen-mediated impairment of T-cell activation and disease induction (43). Myelin oligodendrocyte glycoprotein (MOG)-reactive T cells were generated by immunizing young (2-3-month-old) female wild-type C57BL/6J donors with MOG_35-55_ peptide followed by *ex-vivo* polarization into Th17 phenotype and adoptive transfer into young and aged *Pdgfra-Cre^ERT^*; *RCE* OPC lineage traced mice. Young and aged *Pdgfra-Cre^ERT^*; *RCE* mice that were administered tamoxifen but did not receive adoptive transfer served as naïve controls. For transmission electron microscopy analysis (TEM), two additional experimental lines were used: wild-type C57BL/6J mice that did not receive tamoxifen and OPC Cre combined with the *Tau^mGFP^* reporter (*Pdgfra-Cre^ERT^*; *Tau^mGFP^*) which have been previously used to visualize newly formed myelin sheaths (43–45). *Pdgfra-Cre^ERT^*; *RCE* OPC lineage traced mice were sacrificed at 9-10 days post-adoptive transfer, corresponding to peak disease (Figure 1B) for immunohistochemistry analysis of spinal cord. Wild-type and *Pdgfra-Cre^ERT^*; *Tau^mGFP^*OPC remyelination reporter mice were sacrificed at 10 days post-adoptive transfer for TEM. Extension of the analysis timepoint in aged mice beyond the acute peak disease time-point was not possible without significant loss of aged animals as one third of aged mice reached humane end-point criteria at day 9 (Figure 1C, score ≥4, no movement around cage, inability to feed). Aged adoptive transfer mice had significantly higher experimental autoimmune encephalomyelitis (EAE) clinical scores relative to young adoptive transfer mice (Figure 1B) and faster onset to quadriplegia (Figure 1C). Increased disease severity in aged Th17 adoptive transfer recipients in other studies is not ameliorated by young bone marrow transplantation indicating that the aged host CNS environment drives an exacerbated disease course (46).

**Figure 1.**
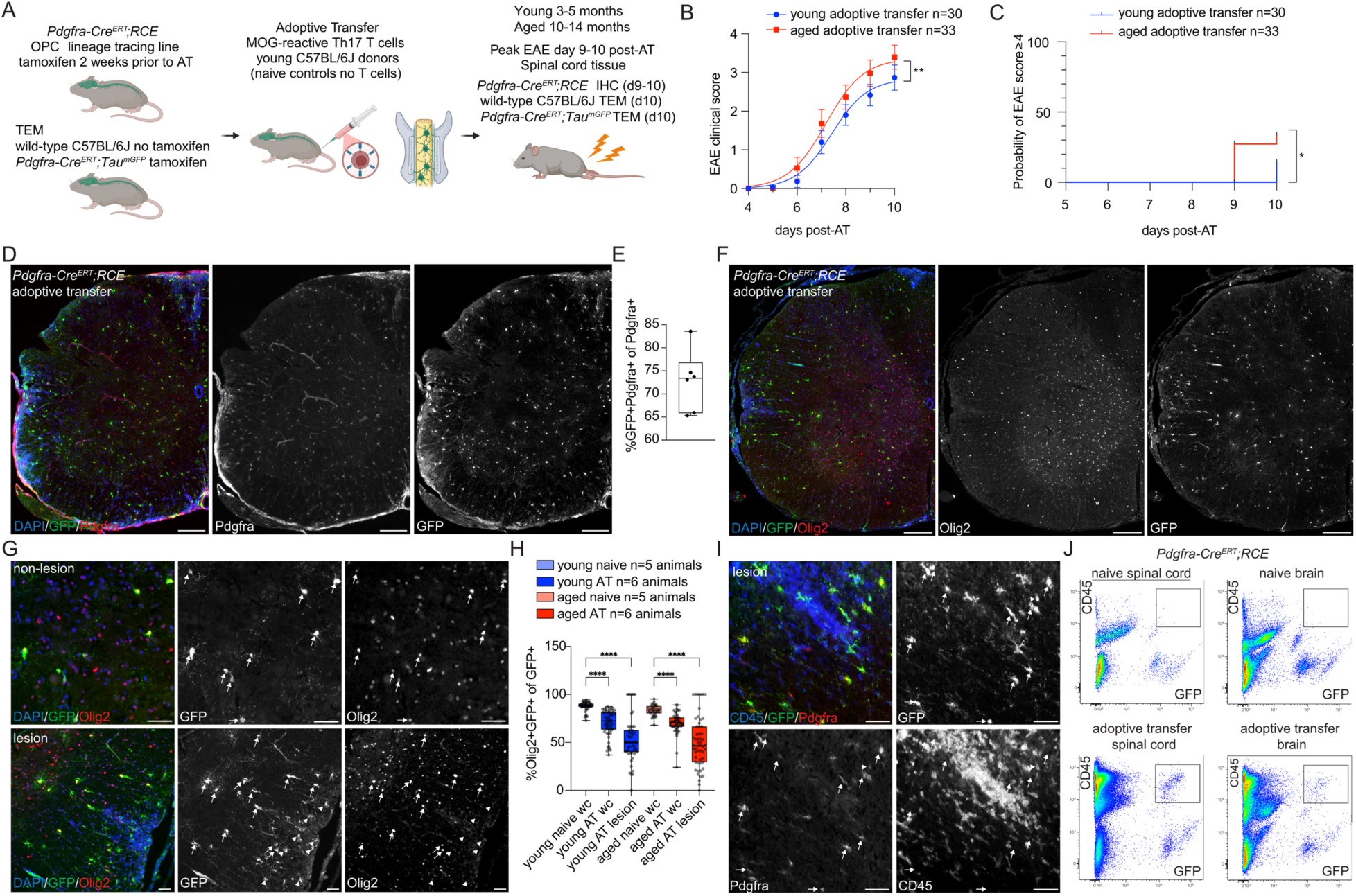
OPC lineage tracing strategy in young and aged Th17 adoptive transfer **(A).** Experimental strategy for MOG-reactive Th17 adoptive transfer (AT) in young and aged *Pdgfra-Cre^ERT^;RCE* OPC lineage traced mice (for immunohistochemistry), wild-type C57BL/6J (for TEM), and *Pdgfra-Cre^ERT^;Tau^mGFP^*OPC remyelination mice (for TEM immunogold). No adoptive transfer young and aged *Pdgfra-Cre^ERT^;RCE* OPC lineage traced mice (for immunohistochemistry) served as naïve controls. Created in BioRender. **(B).** EAE clinical scoring of all young and aged adoptive transfer animals. Aged adoptive transfer animals have significantly higher EAE clinical scores compared to young animals. All young and aged experimental animals pooled. Two-way repeated measures ANOVA, age ** p≤0.001. Non-linear regression sigmoidal variable slope lines, error bars 95% CI. (**C)**. Kaplan-Meier survival analysis of probability of EAE score of 4 or greater (quadriplegia, euthanasia endpoint) in young and aged adoptive transfer with significant difference in time to score 4 endpoint. Logrank (Mantel-Cox) test, * p≤0.05. (**D)**. Representative images of *Pdgfra-Cre^ERT^;RCE* Th17 adoptive transfer spinal cord with lineage-traced OPCs labeled with GFP antibody and Pdgfra. (**E**). Quantification of OPC recombination efficiency in naïve young and aged *Pdgfra-Cre^ERT^;RCE* by percentage of GFP+Pdgfra+ of total Pdgfra+ cells in whole spinal cord section. Data points mean individual animals, n=6 animals. (**F)**. Representative images of *Pdgfra-Cre^ERT^;RCE* Th17 adoptive transfer spinal cord co-labeled with pan-oligodendroglial marker Olig2. (**G)**. Lineage traced GFP+ OPCs express Olig2 (arrows) in spinal cord non-lesion and lesion areas. GFP+Olig2-cells (arrowheads) are present particularly within lesions. (**H)**. Quantification of the percentage of lineage traced GFP+Olig2+ of total GFP+ cells in whole cord (wc) and lesions. Percentage of GFP+ cells that are oligodendroglia (Olig2+) are significantly lower in adoptive transfer wc and lesions compared to naïve wc in both young and aged animals. Median shift Bootstrap test, ****p<0.0001. (**I).** Representative image of lineage-traced *Pdgfra-Cre^ERT^;RCE* adoptive transfer spinal cord lesion co-labeled with GFP, Pdgfra and CD45. GFP+Pdgfra+CD45- OPCs (arrows) and GFP+Pdgfra+CD45+ cells (arrowhead) are present within lesions. (**J)**. Flow cytometry of lineage traced *Pdgfra-Cre^ERT^;RCE* naïve and adoptive transfer brain and spinal cord with GFP+CD45+ cells (box) in adoptive transfer tissue. Box and whiskers plots: box 25 to 75 percentile, whiskers min to max, median line, show all datapoints. Scale bars: D,F 200μm; G,I 50μm.

### Pdgfra-Cre lines lineage trace hematopoietic cells in CNS inflammatory models

Lineage traced *Pdgfra-Cre^ERT^;RCE* mice demonstrated high recombination efficiency and GFP reporter induction in OPCs (Figure 1D) with ∼70% of Pdgfra-positive cells expressing GFP (Figure 1E). Lineage traced GFP reporter-positive cells demonstrated expression of pan-oligodendroglial marker Olig2 (Figure 1F). However, this OPC Cre *Pdgfra-Cre^ERT^* line consistently demonstrated non-oligodendroglial lineage traced cells (GFP+/Olig2-) in adoptive transfer spinal cord particularly within lesions (Figure 1G-H). GFP reporter positive cells were positive for hematopoietic cell marker CD45 and Pdgfra (Figure 1I). Flow cytometry confirmed the presence of a GFP+CD45+ cell population in adoptive transfer CNS tissue (Figure 1J), indicating that peripheral immune cells that traffic into the CNS undergo recombination. These findings highlight the need for rigorous validation of Cre drivers with a lineage tracing approach when using CNS inflammatory disease models. Given the presence of GFP reporter positive non-oligodendroglial cells in adoptive transfer, subsequent quantitative analysis of lineage traced OPCs required inclusion of oligodendroglial restricted markers.

### Young and aged lineage traced OPCs are enriched within adoptive transfer lesions

In the Th17 adoptive transfer model, immune infiltrates are concentrated at leptomeningeal and perivascular spaces of the spinal cord and hindbrain (47–48). To characterize the impact of aging on OPC density and differentiation in response to T cell adoptive transfer, transverse spinal cord cross-sections from young and aged adoptive transfer and naïve animals were stained for lineage traced OPCs with GFP and pan-oligodendroglial marker Olig2 (Figure 2A). Lineage traced OPCs were quantified in total spinal cord cross-sections, lesions (determined by DAPI hypercellularity), and non-lesional white matter. For initial quantification all lesions present in one spinal cord cross-section were grouped together for the total lesional density (Figure 2B). To capture the differences across lesions within a single spinal cord section, individual lesions were quantified separately for downstream analyses (Figure 2C-F, H). Aged lineage traced OPC (GFP+Olig2+) density was significantly lower in adoptive transfer whole cord compared to young, however aged OPCs were present at similar densities in adoptive transfer compared to aged naïve whole cord (Figure 2B) reflecting lower baseline OPC densities with aging. Lineage traced OPC (GFP+Olig2+) density was not significantly different between aged and young lesions when pooling all lesions across a spinal cord section and not accounting for differentiation status (Figure 2B).

**Figure 2.**
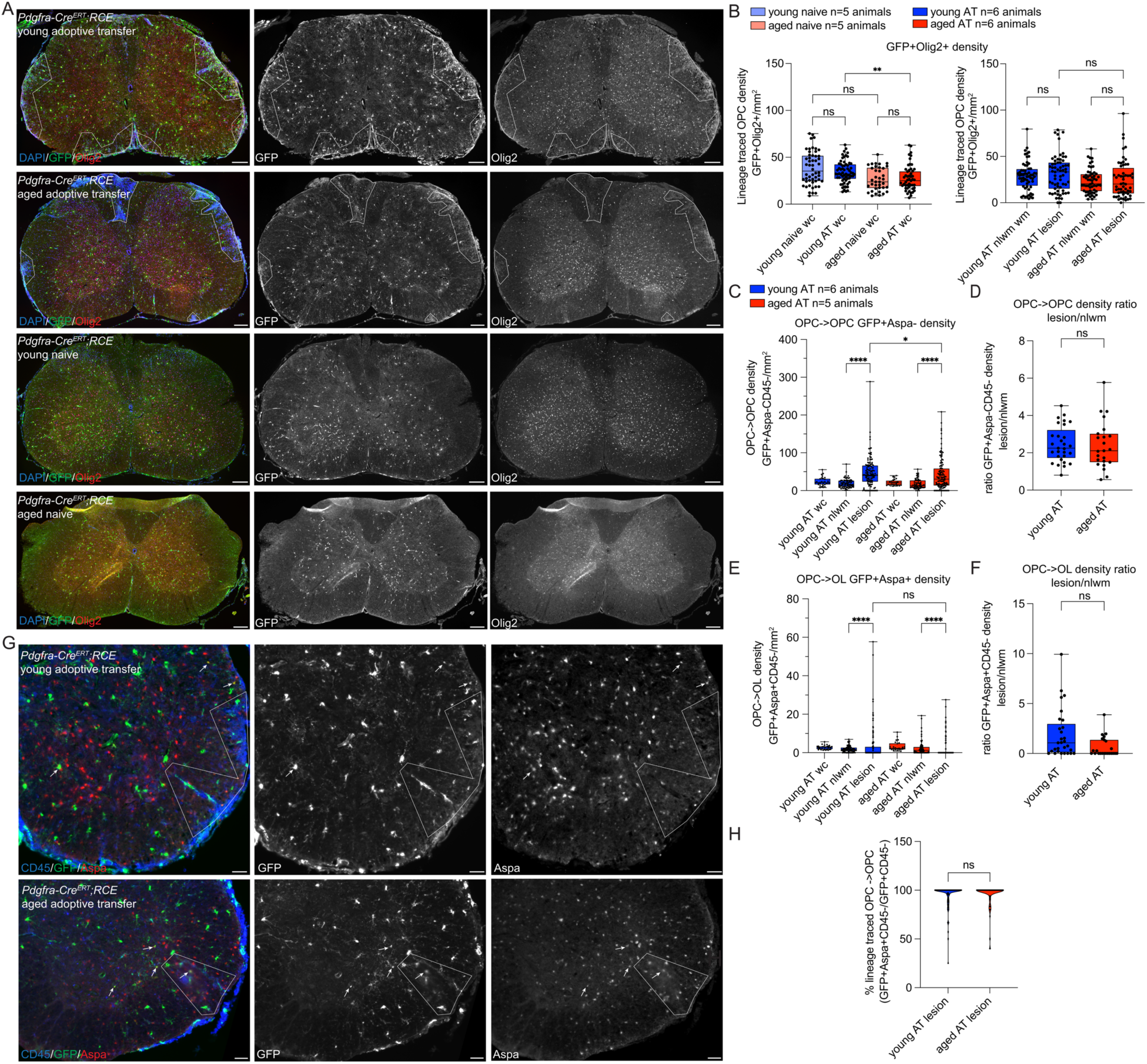
Young and aged lineage traced OPCs are enriched within adoptive transfer lesions **(A).** Representative images of young and aged lineage traced OPC (*Pdgfra-Cre^ERT^;RCE*) naïve and adoptive transfer spinal cord cross-sections stained for pan-oligodendroglial marker (Olig2), lineage traced OPCs (GFP) and DAPI (lesions outlined). (**B)**. Quantification of lineage traced OPC (Olig2+GFP+) densities in whole cord (wc), adoptive transfer (AT) non-lesion white matter (nlwm) and adoptive transfer lesions. Lesion white matter density datapoints are means of all lesions in an individual spinal cord section. Median shift Bootstrap test. (**C)**. Quantification of density of lineage traced OPCs that remained OPCs (OPC->OPC, GFP+Aspa-CD45-) in young and aged adoptive transfer wc, nlwm, and individual lesions. OPC->OPC density is significantly higher in lesions compared to nlwm in young and aged AT. OPC->OPC density is significantly lower in aged compared to young AT lesions. Median shift Bootstrap test. (**D)**. Quantification of the ratio of mean OPC->OPC density in lesions compared to nlwm across a whole spinal cord section. OPC->OPC density is enriched in lesions compared to surrounding nlwm in both young and aged AT and lesion/nlwm ratios are not significantly different between ages. Median shift Bootstrap test. (**E)**. Quantification of density of lineage traced OPCs that differentiated into OLs (OPC->OL, GFP+Aspa+CD45-) in young and aged adoptive transfer wc, nlwm, and individual lesions. Differentiated OPCs OPC->OL density is significantly higher in lesions compared to nlwm in young and aged AT. Median shift Bootstrap test. (**F)**. Quantification of the ratio of mean OPC->OL density in lesions compared to nlwm across a whole spinal cord section. OPC->OL lesion/nlwm ratios are not significantly different between ages. Median shift Bootstrap test. (**G)**. Representative images of young and aged *Pdgfra-Cre^ERT^;RCE* adoptive transfer spinal cord stained for GFP, Aspa, and CD45. OPCs that have differentiated (GFP+Aspa+) are indicated with arrows. Lesions outlined by presence of DAPI hypercellularity (not shown) and CD45 infiltrate. (**H)**. Quantification of percentage of lineage traced OPCs that remain OPCs (GFP+Aspa-CD45- of GFP+CD45-) within young and aged individual lesions. Median shift Bootstrap test. Box and whiskers plots: box 25 to 75 percentile, whiskers min to max, median line, show all datapoints. p values: ns p>0.05, * p≤0.05, **p≤0.01, ***p≤0.001, ****p<0.0001. Scale bars: A, 200μm; F 50μm.

To better characterize the fate and differentiation status of lineage traced OPCs, spinal cord sections were co-labeled with GFP, mature oligodendrocyte marker (Aspa), and hematopoietic marker (CD45) followed by quantification of all GFP+CD45- cells in spinal cord cross-sections (Figure 2C-H). In both young and aged animals, lineage traced OPCs that remained OPCs (OPC->OPC, GFP+Aspa-CD45-) were significantly increased within lesions compared to non-lesional white matter (Figure 2C). OPC->OPC density was significantly lower in aged compared to young lesions (Figure 2C). To determine the enrichment of OPCs within lesions compared to surrounding non-lesional white matter, a ratio of mean lesional density over non-lesional white matter (nlwm) density was calculated for each spinal cord cross-section (Figure 2D). Lineage traced OPCs that remained OPCs (GFP+Aspa-CD45-) had a greater than 2-fold enrichment in both young and aged lesions compared to surrounding nlwm suggesting recruitment, proliferation, and/or retention of OPCs within the lesional environment.

### Young and aged OPCs demonstrate the ability to differentiate in acute adoptive transfer

The density of OPCs increased in lesions of young and aged animals, suggesting that despite the more severe disease observed in aged animals, OPCs retain the capacity to respond to demyelination and potentially differentiate. A small minority of lineage traced OPCs underwent differentiation and expressed Aspa (GFP+Aspa+CD45-) in both young and aged adoptive transfer spinal cord (Figure 2E). Compared to non-lesion white matter, differentiated OPCs were significantly higher in young lesions and significantly lower in aged lesions (Figure 2E). Young OPCs may have a higher or faster capacity for differentiation within lesions and/or aged OPCs undergoing differentiation may be more susceptible to cell death in the aged environment. Comparing the ratio of newly differentiated OPCs to surrounding nlwm on an individual spinal cord section level (Figure 2F) did not indicate a significant difference between the ratio of differentiated OLs within young and aged lesions compared to surrounding nlwm (Figure 2F). To determine the percentage of lineage traced OPCs that remained OPCs within lesions, all lineage traced OPCs (GFP+CD45-) were quantified within individual lesions and assessed for Aspa expression. Similar to density analysis, lineage traced OPCs that remained OPCs made up over 95% of lineage traced OPCs within lesions of young and aged animals (Figure 2H). The enrichment of lineage traced OPCs within young and aged lesions and ability of OPCs to differentiate, particularly within young lesions at this peak disease time-point, indicates that OPCs can quickly respond even with ongoing inflammatory activity and high disease severity. Alternative immune-mediated models with remission of clinical disease may offer additional insight into the kinetics of OPC responses after resolution of inflammatory activity.

### Young and aged differentiated oligodendrocytes are lost in adoptive transfer

Adoptive transfer of myelin oligodendrocyte glycoprotein (MOG)-reactive Th17 T cells may result in targeted cytotoxicity to MOG-expressing mature OLs. To determine whether OLs are lost or preserved in young and aged adoptive transfer, spinal cord sections were stained for OL marker Aspa (Figure 3A). High densities of differentiated OLs in spinal cord cross-sections necessitated the development of a semi-automated cell counting method. An ImageJ macro was developed to quantify Aspa-positive cell densities (Figure 3B). Aspa+ OL densities were significantly lower at baseline in aged compared to young naïve spinal cord (Figure 3C). Adoptive transfer resulted in a significant decrease in OL density by ∼1/4 of density relative to naïve controls (Figure 3C) and aged adoptive transfer spinal cord had the highest degree of OL loss. To determine if existing OLs (not derived from OPCs) were selectively lost within lesions we performed manual quantification of OLs in individual lesions and adjacent nlwm with use of GFP/Aspa/CD45 co-labeling (Figure 2G). Existing differentiated OLs (Aspa+GFP-CD45-) were significantly lower in lesions compared to non-lesions in both young and aged animals with the lowest density of OLs within aged lesions (Figure 3D). The ratio of OL lesional to non-lesional density was not significantly different between young and aged adoptive transfer (Figure 3E).

**Figure 3.**
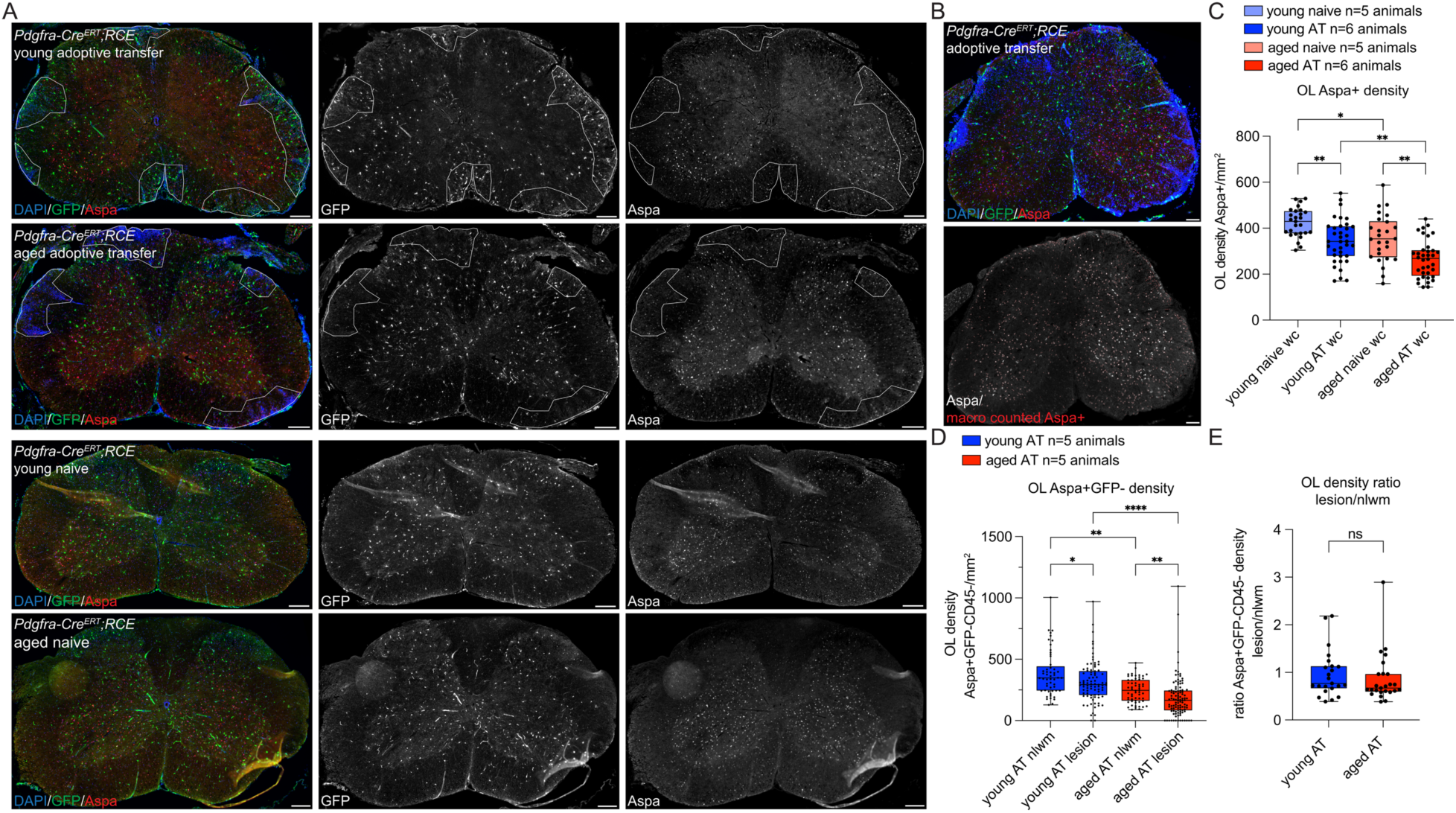
Differentiated oligodendrocytes are reduced in adoptive transfer **(A).** Representative images of spinal cord sections from young and aged lineage traced OPC *Pdgfra-Cre^ERT^;RCE* adoptive transfer and naïve spinal cord sections stained for differentiated oligodendrocyte (OL) marker (Aspa), lineage traced OPCs (GFP) and DAPI (lesion areas outlined). (**B)**. Representative adoptive transfer spinal cord section stained for DAPI/GFP/Aspa and processed with GFP/Aspa ImageJ macro with Aspa-positive macro counted cells outlined in red overlayed on raw grayscale of Aspa staining (bottom panel). (**C)**. Quantification of Aspa+ OL densities with macro in young and aged naïve and adoptive transfer (AT) whole cord (wc) cross-sections. Aspa+ OL density is significantly lower in aged naïve compared to young naïve and in adoptive transfer compared to naïve in both ages. Median shift Bootstrap test. (**D)**. Manual quantification of differentiated Ols (GFP-Aspa+CD45-) in young and aged adoptive transfer non-lesion white matter (nlwm) and individual lesions. OL density is significantly lower in lesions compared to nlwm in both ages. Aged AT OL density is significantly lower than young AT in both regions. Median shift Bootstrap test. (**E)**. Quantification of the ratio of mean OL density in lesions compared to nlwm across a whole spinal cord section. OL lesion/nlwm ratios are not significantly different between ages. Median shift Bootstrap test. Box and whiskers plots: box 25 to 75 percentile, whiskers min to max, median line, show all datapoints. p values: ns p>0.05, * p≤0.05, **p≤0.01, ****p<0.0001. Scale bars: A,B 200μm.

The lower density of aged OLs throughout spinal cord regions may reflect overall higher disease severity in aged animals and/or increased susceptibility of aged OLs to cell death. To investigate the impact of EAE score and CD45 infiltrate on OPC and OL densities within the spinal cord of young and aged adoptive transfer animals we performed a correlation analysis on matched cell counts from individual spinal cord sections. Variables of age, EAE score, percentage of cord occupied by CD45 infiltrate, and density of OPCs (GFP+Aspa-CD45-), OPC->OL (GFP+Aspa+CD45-), and OLs (GFP-Aspa+CD45-) across the whole cord and mean of pooled lesions were compared for each spinal cord section (Supplemental Figure 1A). Strong correlations (r values <-0.5 or >+0.5 and p values of ≤0.001) included the following correlations: positive correlations-age to EAE score, OPCs in cord to OPCs in lesion, OPCs in cord to OPC->OL in cord, and negative correlation-age to OL lesion. EAE score was not significantly correlated with OPC/OL densities. Degree of CD45 infiltrate was not correlated with age, EAE score or OPC/OL densities. While pooling counts from lesions may limit the power of this analysis, it suggests that the range of EAE scores across aged and young animals does not markedly impact OPC/OL densities. Notably, higher levels of OPCs throughout the whole cord was correlated with higher OPCs within lesions and OPCs that differentiated in the cord. Higher age may also correlate with reduced OL lesional densities which was also suggested by our macro and individual lesional quantification (Figure 3C-D).

### Aging increases myelin debris accumulation after adoptive transfer

To determine the impact of aging on the degree of axonal loss, myelin debris and demyelination after adoptive transfer, transverse spinal cord sections from young and aged adoptive transfer mice were stained for neurofilament (NF), degraded myelin basic protein (dMBP), and FluoroMyelin (total myelin) (Figure 4). To quantify axonal loss within lesions an ImageJ macro was developed to quantify neurofilament-positive axonal density (Figure 4A). Axonal density was highly variable across lesions, and no significant differences were observed between young and aged lesions (Figure 4B). The accumulation of myelin debris was assessed by the presence of dMBP staining (Figure 4C) and lesions in aged animals had a significantly higher percentage of white matter with positive dMBP staining (Figure 4D). Higher levels of myelin debris in aged animals may reflect impaired myelin debris clearance by aged macrophages (20). Although myelin debris was higher in aged animals, there was no significant differences in percent of white matter demyelination quantified by FluoroMyelin staining between age groups (Figure 4E-F).

**Figure 4.**
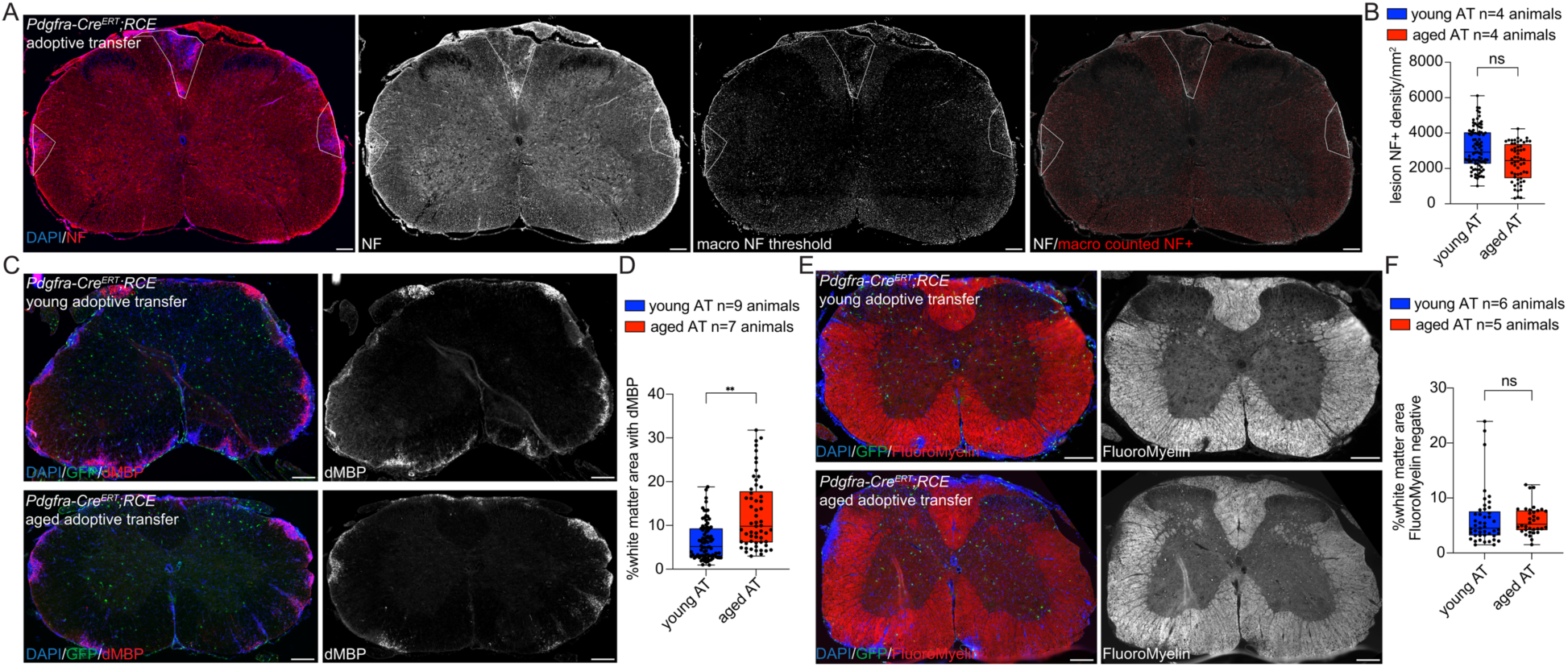
Axonal loss, myelin debris and demyelination in young and aged adoptive transfer **(A).** Representative images of macro quantification of axonal density in *Pdgfra-Cre^ERT^;RCE* adoptive transfer spinal cord section stained for neurofilament-H (NF) and DAPI (left panel), NF only grayscale image (middle left panel), NF macro thresholded image (middle right panel) and NF macro processed image with NF+ axons outlined in red (right panel). Lesions outlined. (**B)**. Quantification of NF axonal density by macro in individual lesions of young and aged adoptive transfer (AT). No significant difference in axonal density in lesions between ages. Median shift Bootstrap test. **(C)**. Representative image of myelin debris/degraded myelin (dMBP) and GFP staining in young and aged *Pdgfra-Cre^ERT^;RCE* adoptive transfer spinal cord sections. (**D)**. Quantification of the percentage of white matter area with dMBP staining in young and aged AT spinal cord with significant increase white matter with myelin debris in aged white matter. Median shift Bootstrap test. (**E)**. Representative images of FluoroMyelin and GFP staining in young and aged adoptive transfer *Pdgfra-Cre^ERT^;RCE* spinal cord sections. White matter regions with DAPI hypercellularity demonstrate absence of FluoroMyelin staining. (**F)**. Quantification of the percentage of demyelinated white matter with no FluoroMyelin staining in young and aged AT spinal cord with no significant difference between ages. Median shift Bootstrap test. Box and whiskers plots: box 25 to 75 percentile, whiskers min to max, median line, show all datapoints. p values: ns p>0.05, **p≤0.01. Scale bars: A,C,E 200μm.

### Remyelination is present to a similar extent in young and aged lesions

In the Th17 adoptive transfer model, areas of demyelination are localized in the peripheral white matter adjacent to leptomeningeal infiltrates indicated by DAPI hypercellularity (Figure 4E) (47–48). To determine if OPCs can remyelinate in adoptive transfer, we crossed OPC inducible Cre line *Pdgfra-Cre^ERT^* to inducible membrane targeted GFP within the microtubule associated protein tau locus (*Tau^mGFP^*) which has been used to visualize newly generated myelin sheaths (43–45). Newly generated OL GFP+ myelin sheaths were not seen in naïve spinal cord (Figure 5A). In adoptive transfer, GFP+ processes surrounding axons and co-labeled with MBP were visualized at the edge of lesions (Figure 5B) suggesting the presence of remyelination from newly formed OLs. Immunohistochemistry of a young and aged adoptive transfer cohort of remyelination reporter mice (*Pdgfra-Cre^ERT^;Tau^mGFP^*) was performed however quantification of remyelination was not pursued due to GFP reporter background expression in OPC processes in the parenchyma (Figure 5B).

**Figure 5.**
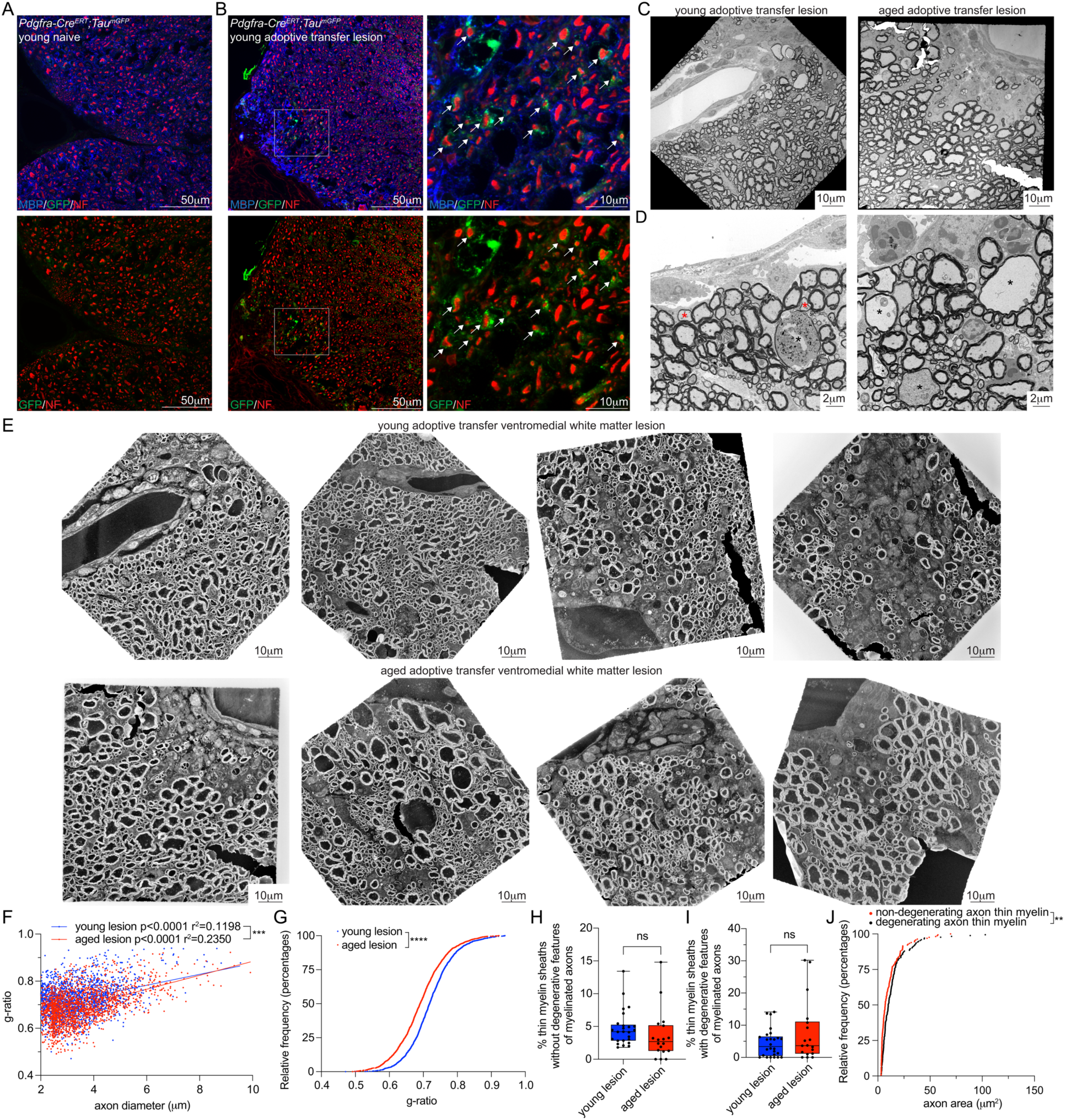
Remyelination in young and aged adoptive transfer lesions **(A).** Representative image of lumbar ventrolateral spinal cord in lineage traced OPC Cre line crossed to membrane GFP reporter (*Pdgfra-Cre^ERT^;Tau^mGFP^*) for labeling remyelinating myelin sheaths from OPCs. Staining for myelin (MBP), lineage traced OPC processes (GFP) and axons (NF) with no GFP+ myelin sheaths in naïve/non-adoptive transfer. **(B)**. Representative image of young *Pdgfra-Cre^ERT^;Tau^mGFP^*ventral spinal cord 10 days post-adoptive transfer with lineage traced OPC GFP+ processes wrapping NF+ axons (arrows) at the lesion edge, box indicates region of higher magnification. (**C)**. Representative transmission electron microscopy (TEM) images of ventromedial white matter lesions in young and aged adoptive transfer. (**D)**. Higher magnification TEM images of a young adoptive transfer lesion with thin myelin sheaths around degenerating axons with electron dense material or myelin infoldings (black asterisks) and thin myelin sheaths around axons without signs of degeneration (red asterisks). (**E)**. Representative TEM inverted images of young and aged adoptive transfer lesions to highlight myelin sheaths. (**F)**. Quantification of g-ratios of axons ≥2μm diameter in young and aged adoptive transfer lesions. Simple linear regression analysis. (**G)**. Cumulative relative frequency distribution of g-ratios indicating a significantly higher proportion of lower g-ratios (thicker myelin sheaths) in aged lesions. Kolmogorov-Shirnov test. (**H)**. Quantification of percent of thin myelin sheaths (g-ratio ≥0.8) without degenerative features of total myelinated axons with diameter ≥2μm within a lesion. No significant difference between young and aged lesions. Median shift Bootstrap test. (**I)**. Quantification of percent of thin myelin sheaths (g-ratio ≥0.8) with degenerative features of total myelinated axons with diameter ≥2μm within a lesion. No significant difference between young and aged lesions. Median shift Bootstrap test. (**J)**. Cumulative relative frequency distribution of axonal areas of thin myelin sheaths with degenerative features (black) or no degenerative features (gray) with a significantly higher proportion of large axons in axons with degenerative features. Kolmogorov-Shirnov test. F-G, n=4 young animals with 31 lesions, n=6 aged animals with 36 lesions. H-J, n=4 young animals with 25 lesions, n=6 aged animals with 18 lesions. Box and whiskers plots: box 25 to 75 percentile, whiskers min to max, median line, show all datapoints. p value: ns, p>0.05, **p≤0.01, ***p≤0.001, ****p<0.0001.

To more reliably assess remyelination transmission electron microscopy (TEM) was performed on wild-type C57BL/6J young and aged ventromedial white matter lesions at 10 days post-adoptive transfer. Longer time-point analysis was not possible in aged animals due to increased morbidity in this model (Figure 1C). Ventromedial white matter was chosen based on the high rate of lesions in this region, ability to identify landmarks with the presence of a ventral vein, and large diameter axons (49) that could facilitate the ability to detect thinner remyelinated axons. Toxin-mediated demyelinating models have demonstrated that remyelinated axons can be identified by thinner myelin sheaths (50–52), however thin myelin sheaths can also be a sign of degenerating axons both in toxin-mediated and immune-mediated models (53). Perivascular immune infiltrates lining areas of demyelination were identified in all ventromedial white matter sections imaged as well as high concentration of oligodendrocytes within demyelinated areas (Figure 5C). Thin myelin sheaths surrounding axons with degenerative features (electron dense material, myelin blebbing, myelin infoldings, myelin unraveling, high mitochondrial concentration) and axons without these degenerative features were present within lesions (Figure 5D). TEM lesion images at 1150x magnification were used for quantification of young and aged lesions with inverted images facilitating the visualization of myelin sheaths and axonal components (Figure 5E). G-ratio quantification was performed on 1150x magnification images of young and aged ventromedial white matter lesions (non-inverted) using MyelTracer (54) combined with manual outlining for axons with a diameter ≥2μm. Axons with thin myelin sheaths were noted to have g-ratios of 0.8 or higher. Axons with g-ratio ≥ 0.8 were present in both age groups. G-ratio slopes were significantly different between young and aged lesions (Figure 5F). Relative frequency distribution analysis of g-ratios demonstrated a significantly higher frequency of lower g-ratios (thicker myelin) in aged lesions (Figure 5G), indicating that the difference in g-ratio slope was driven by thicker myelin sheaths in aged compared to young animals. This finding is consistent with previous studies of human (55) and mouse spinal cord tracts (56) which have demonstrated thicker myelin with aging. With thin myelin sheaths surrounding both degenerating axons and axons without degenerative components (Figure 5D) we further analyzed axons with thin myelin sheaths (g-ratio ≥0.8) for the presence or absence of degenerative features. Thinly myelinated axons without degenerative features, likely representing remyelinated axons, were present in similar proportions in aged and young adoptive transfer lesions (Figure 5H). Thinly myelinated axons with degenerative features were also present at similar proportions in young and aged lesions (Figure 5I). Relative frequency distribution analysis of axonal cross-sectional area of thinly myelinated axons classified as degenerating compared to non-degenerating indicated that degenerating axons had a significantly higher frequency of large axonal areas (Figure 5J) as would be expected for degenerating axons.

We acknowledge that thin myelin sheaths may represent remyelinated axons or axons in the process of demyelination and our time-point analysis at 10 days-post adoptive transfer would require fast kinetics for OPC differentiation and remyelination. Young and aged OPC remyelination reporter mice *Pdgfra-Cre^ERT^;Tau^mGFP^* that received adoptive transfer were processed for TEM with immunogold staining for GFP to attempt to further validate OPC remyelination. However, due to GFP reporter expression in OPC processes in the parenchyma (Figure 5B) and anti-GFP immunogold deposition in similar pattern, analysis of TEM/immunogold was not further pursued. Despite these limitations, remyelination appears similar between age groups, even in the presence of a robust inflammatory infiltrate and severe disease model.

## Discussion

OPCs are dynamic cells with the ability to self-renew, migrate, and differentiate into mature, myelinating oligodendrocytes. Although the response of OPCs to demyelination has been investigated in toxin-mediated models and genetic ablation strategies (5), the fate of OPCs after exposure to a robust auto-immune inflammatory insult remains poorly understood. Biological aging influences multiple sclerosis disease activity and progression (13) and the response of OPCs in an aged environment is particularly relevant for a chronic CNS autoimmune disease that affects individuals at all stages of life. Physiological aging in the absence of a disease model is associated with a decreased rate of OPC proliferation (14,16) and differentiation into mature OLs (14). Aged OPCs express higher levels of the cell cycle inhibitors p16 and p21, which may reflect age-associated senescence (57). In response to toxin-mediated demyelination insults, aged OPCs have slower proliferation, recruitment, and differentiation kinetics; however, remyelination in aged animals eventually catches up to young animals (17–19).

To determine whether aging impacts the response of OPCs to a strong adaptive immune response akin to an acute MS attack, we performed lineage tracing of OPCs in young adult and aged spinal cord after adoptive transfer of myelin reactive Th17 polarized T cells. Both aged and young OPCs respond to an acute inflammatory demyelination event with a proportionally similar increase in lesional OPC density. The highest densities of lineage traced OPCs in both aged and young animals were within lesions, suggesting that compensatory mechanisms allow for recruitment, survival and persistence of OPCs within a lesion environment. Both young and aged OPCs can differentiate into mature OLs, a response that is less robust in aged OPCs. While existing differentiated OLs sustain damage after adoptive transfer, the magnitude of aged OL loss is more severe. Although the presence of ongoing axonal degeneration in adoptive transfer and other models can complicate the interpretation of thin myelin sheaths by TEM analysis (53), we quantified thin myelin sheaths by TEM and found a similar proportion of non-degenerative and likely remyelinated axons in lesions of young and aged animals. With TEM we were unable to distinguish whether remyelination occurs from newly generated oligodendrocytes versus existing differentiated oligodendrocytes, both of which can occur in toxin-mediated models (4). Further lineage tracing studies using other transgenic membrane targeted reporter lines may offer additional insight into the origin of remyelinating oligodendroglia in immune-mediated models.

Previous studies combining genetic CNS interferon gamma production with toxin-mediated demyelination indicate that OPCs may by targets of CD8 T cell mediated killing (33). Our findings in the Th17 adoptive transfer model indicate that an inflammatory lesion environment stimulates OPCs responses and both young and aged OPCs may have compensatory strategies that allow them to survive and differentiate despite the presence of T cells. Interferon-response pathways are upregulated in disease-associated oligodendroglia (25–32) and with physiological aging (16, 58) which may be a result of CD8 T cell accumulation within the aged CNS (58). Oligodendrocytes upregulate MHC class I (59) and immune checkpoint inhibitor programmed death-ligand 1 (PD-L1) (60) in CNS viral models. Upregulation of interferon-response pathway in young and aged OPCs may in fact be beneficial for oligodendroglial survival allowing OPCs and OLs to evade the immune attack and dampen immune signaling. Further studies are needed to investigate the role of disease-associated oligodendroglia in inflammatory models to determine whether immunoregulatory pathways in oligodendroglia are beneficial or inhibitory to oligodendroglial function.

This study demonstrates that oligodendroglia in young and aged animals respond similarly to an acute auto-immune mediated demyelinating attack mediated by MOG-reactive Th17 T cells. The ability of both young and aged OPCs to respond to a severe autoimmune attack and preservation of OPCs within lesions suggests that remyelination and repair strategies that target oligodendroglial survival mechanisms may be promising strategy for people of all ages with MS.

## Methods

### Sex as a biological variable

Both male and female mice transgenic and wild-type mice were used as adoptive transfer recipients with a roughly equal even distribution between female and male mice in young and aged cohorts. Due to the research strategy involving use of transgenic lines, timing of adoptive transfer cohorts and need for aged cohorts, all available positive transgenic mice were used regardless of sex and the experiments were not sufficiently powered to assess differences between sexes. Wild-type mice used for immunization for donor T cells were female only to avoid potential sex-based T cell differences.

### Animal care and use

Young (P87-P148, 3-5-month-old) and aged (P295-P427,10-14-month-old) *Pdgfra-Cre^ERT^* hemizygote +/- (Jackson Laboratory 018280); *RCE- Rosa26^flstopflEGFP^* hemizygote +/- (Jackson Laboratory 032037-JAX, MMRRC stock #32027). Young (P131, 4-month-old) and aged (P366-P426, 12-14-month-old) C57BL/6J (Jackson Laboratory 000664) wild-type mice and *Pdgfra-Cre^ERT^* hemizygote +/- (Jackson Laboratory 018280); *Tau^mGFP^* hemizygote +/- (Jackson Laboratory 021162) were used for transmission electron microscopy experiments. Age indicated is animal age at sacrifice. Female and male mice were assigned to experiments based on availability. Mice were maintained on a 12-hour light/dark cycle, housed in cages of five or less, and provided food and water ad libitum. Adoptive transfer mice were monitored daily between days 4-10 post-adoptive transfer for clinical score and weight. All animal experiments were conducted in accordance with the National Institute of Health Guide for the Care and Use of Laboratory Animals.

### Adoptive transfer/Experimental autoimmune encephalomyelitis (EAE)

Transgenic mice *Pdgfra-Cre^ERT^*; *RCE* and *Pdgfra-Cre^ERT^;Tau^mGFP^*were administered tamoxifen (Sigma T5648) intraperitoneally (100-150ul of 10mg/ml stock reconstituted in corn oil for working dose of 75mg/kg/day) daily for 5 days to induce recombination starting two weeks prior to adoptive transfer. Wild-type C57BL/6J mice used for transmission electron microscopy (TEM) did not receive tamoxifen. T cells for adoptive transfer were generated from wild-type C57BL/6J (Jackson Laboratory 000664) 2-3-month-old female mice which were anesthetized with intraperitoneal injection of ketamine (90mg/kg)/ xylazine (10mg/kg) and immunized with four subcutaneous flank injections with a total of 0.2-0.4mg of myelin oligodendrocyte glycoprotein (MOG)_35-55_ peptide (1mg/ml, biosynthesis) resuspended in Complete Freund’s Adjuvant (CFA) with concentration of 5mg/ml heat killed *M. tuberculosis*. After 10 days immunized wild-type mice were euthanized with CO_2_ and spleen and lymph nodes were dissected and placed on ice in EAE media: RPMI (ThermoFisher 21870084), 10% FBS (GeminiBio 100-106-500), P/S (Fisher 15-140-148), 1x GlutaMAX (Fisher 35-050-061), 1x NEAA (ThermoFisher 11140050), 1mM sodium pyruvate (Fisher 11-360-070), 55nM beta-mercaptoethanol (ThermoFisher 21985023). Spleens and lymph nodes were mashed across filter, washed with EAE media, and centrifuged at 800g 5min. Spleens were resuspended in RBC lysis buffer (ThermoFisher 00-4300-54) for 2 minutes at room temperature followed by quenching with EAE media, centrifuged at 800g 5min, and washed with EAE media. Spleen and lymph nodes were resuspended in EAE media with MOG_35-55_ peptide (50μg/ml, Biosynthesis), recombinant mouse IL-23 (8ng/ml, R&D 1887-ML), recombinant mouse IL-1a (10ng/ml, ThermoFisher 211-11A), anti-IFNg (10mg/ml, Bio X Cell BE0055) at volume of 5×10^6^ cells/ml in T75 flasks. Cells suspensions were incubated at 37°C in 5% CO_2_ incubator for 3.5 days. After *ex-vivo* Th17 polarization cells were collected into conical tubes, centrifuged 800g 5min, filtered and washed twice with MACs buffer (0.5% BSA and 2mM EDTA in PBS). CD4 T cell selection was performed with mouse CD4 (L3T4) microbeads (Miltenyi Biotec 130-117-043) with LS column per Miltenyi protocol. Eluted CD4+ cells were washed in MACs buffer twice and 4×10^6^ cells were injected intraperitoneally into transgenic mice (tamoxifen administered) and wild-type mice (no tamoxifen). For naïve/non-adoptive transfer animals, littermates of transgenic mice that received tamoxifen were separated into naïve cages. Mice were monitored daily starting on day 4 post-adoptive transfer with weights and EAE clinical scoring scale. EAE scoring scale: 0.5 reduced tail tone; 1 complete loss of tail tone; 1.5 slight gait imbalance; 2 gait imbalance with footfalls or spinning/ataxia; 2.5 hindlimb foot drag able to move above hip; 3 partial to complete hindlimb paralysis, not able to move either hindlimb above hip, moving around cage with forelimbs; 3.5 complete hindlimb paralysis, hindlimbs held to one size, moving around cage with forelimbs; 4 complete hindlimb paralysis, hindlimbs held to one size, no movement around cage but alert; 4.5 moribund not alert; 5 death. Mice were sacrificed at days 9-10 post-adoptive transfer with most severe mice sacrificed on day 9 time-point to avoid death and following end-point euthanasia protocols (euthanasia at score 4 or greater within 24 hours or sustained weight loss ≥30% for 3 days). All mice in this study were between clinical scores of 2-4.5 at day of sacrifice and a total of 5 separate cohorts of adoptive transfer were performed. EAE clinical score graphs from mice used for immunohistochemistry and transmission electron microscopy including several additional cohorts that were used to optimize TEM fixation protocol were combined for EAE clinical score graphs. There was no significant difference between mice that received tamoxifen and those that did not. Animals that were sacrificed due to endpoint criteria or reached score 5 (death) were included as data-points with score 5 on subsequent days on EAE clinical scoring graphs.

### Immunohistochemistry

Mice were anesthetized with intraperitoneal injection of ketamine (90mg/kg)/ xylazine (10mg/kg) and perfused transcardially with 15ml of 0.1M phosphate-buffered saline (PBS) followed by cold 4% paraformaldehyde (PFA) in PBS (ThermoScientific J19943.K2). Spinal cords were dissected out of the spinal column and post-fixed in 4% PFA/PBS at 4°C overnight then cryoprotected in 30% sucrose/1xPBS at 4°C for at least 2 days. Prior to embedding two transverse cuts were performed (at the end of the cervical enlargement and beginning of lumbar enlargement) to separate the spinal cord into cervical, thoracic and lumbar segments. Spinal cord segments were embedded in Tissue-Tek OCT compound (Sakura) with a cross-section of the beginning of each spinal cord segment facing the cutting edge of the block. Spinal cord blocks were sectioned at 15μm thickness with Leica CM1950 cryostat onto Superfrost Plus microscope slides (Fisher 12-550-15) with each 8-10 series of slides having four sets of cervical/thoracic/lumbar transverse sections (12 total sections) representing ∼600mm distance from first to fourth section in each region to obtain representative analysis across each spinal cord segment on one slide. Slides were stored at −80°C until use. For immunostaining slides were permeabilized with 1xPBS +0.1% Triton-X100 (PBST) for 10 minutes followed by outlining of sections with PAP pen (ThermoFisher 008899). To prevent non-specific binding, 5% normal goat serum (JacksonImmuno 005-000-121) in PBST was added to sections for 1 hour at room temperature followed by addition of primary antibodies in same blocking solution overnight at 4°C. Slides were washed in PBST three times 5 minutes each followed by incubation in secondary antibodies in blocking buffer for 1 hour at room temperature. Slides were washed with PBST twice for 5 minutes each, once with PBS for 5 minutes then mounted with ProlongGold antifade (ThermoFisher P36935). For FluoroMyelin co-stained slides, FluoroMyelin Red (Invitrogen F34652) was added at 1:300 dilution in PBS after last wash and incubated for 20 minutes followed by PBS wash and mounting. The following primary antibodies were used: chicken anti-GFP (Abcam ab13970, 1:1000), rabbit anti-Olig2 (MilliporeSigma ab9610, 1:1000), rabbit anti-Aspa (MilliporeSigma ABN1698, 1:250), rat anti-CD45 (MilliporeSigma 05-1416, 1:500), rabbit anti-degraded myelin basic protein dMBP (MilliporeSigma ab5864, 1:1000), rabbit anti-Pdgfra (CellSignaling 3174, 1:500), mouse anti-neurofilament heavy NF-H (EnCor MCA-AH1), rat ant-MBP (Biorad MCA409S, 1:1000), mouse anti-neurofilament medium NF-M (Invitrogen 130700, 1:1000, for remyelination reporter staining). The following secondary antibodies were used all purchased from Invitrogen and used at dilution of 1:500: Alexa Fluor 488 goat anti-chicken IgY (H+L) (A-11039), Alexa Fluor 594 goat anti-rat IgG (H+L) (A-11007), Alexa Fluor 647 goat anti-rat IgG (H+L) (A-21247), Alexa Fluor 647 goat anti-rabbit IgG (H+L) (A-21245), Alexa Fluor 647 goat anti-mouse IgG (H+L) (A-21236), and Alexa Fluor 594 goat anti-mouse IgG (H+L) (A-11032).

### Immunohistochemistry imaging and analysis

Images of immunohistochemistry-stained spinal cord sections were acquired using Olympus IX83 inverted microscope with Zeiss ZEN microscopy software. Tiled images of transverse spinal cord sections at a resolution of 650nm/pixel (10x objective) were obtained for GFP/Olig2, GFP/Aspa, and dMBP stains. Higher resolution tiled images at 325nm/pixel (20x objective) were taken for GFP/Aspa/CD45, neurofilament and FluoroMyelin stains. For manual-counting, image stacks were analyzed in ImageJ/Fiji using channels tool, cell counter and ROI manager. For macro-analyzed images (DAPI/GFP/Aspa and DAPI/GFP/NF stains), images were exported as non-compressed TIFF grayscale stack of channels. Macros were developed using a combination of background subtraction, moments thresholding and analyzing particle ImageJ/Fiji functions to identify positive cells. In refinement of macros for individual stains, quality assessment images with overlay of counted objects were analyzed for accuracy and parameters were adjusted for each macro to obtain the highest accuracy of cell detection. After importing the spinal cord section image stack, the macros prompt the user to outline regions of interest (ROI)- whole cord, gray matter and lesions. Lesions were identified and outlined by the presence of DAPI hypercellularity which correlated with CD45 infiltrate. Non-lesion white matter counts were generated by subtracting whole cord counts from gray matter and total lesion counts, this was due to the ease in outlining gray matter during the macro execution compared to total non-lesion white matter. For Aspa macro counts total lesions within each spinal cord section were outlined as one lesion. In NF macro and DAPI/GFP/Aspa/CD45 manual counts individual lesions were outlined separately to increase power of analysis given potential variability across lesions. For neurofilament quantification, DAPI/GFP/NF 20x tiled images were analyzed with NF macro allowing users to outline lesions based on DAPI hypercellularity and NF positive axonal density was calculated for individual lesions. For dMBP quantification DAPI/GFP/dMBP 10x tiled image stacks were quantified in ImageJ/Fiji by outlining total white matter area and white matter with positive dMBP staining for each spinal cord section. For FluoroMyelin quantification, DAPI/GFP/Fluoromyelin 20x tiled image stacks were quantified by outlining total white matter and demyelinated white matter without Fluoromyelin. To generate non-biased transverse spinal cord cross-section counts throughout the cervical, thoracic and lumbar spinal cord, all spinal cord sections without significant folds or other tissue processing artifacts were imaged. Adoptive transfer sections without lesions were not quantified to focus on spinal cord regions with inflammatory infiltrate. Sections quantified per animal: GFP/Olig2 6-16, GFP/Aspa/CD45 hand counts 3-6, GFP/Aspa macro 6, GFP/NF macro 3-8, GFP/FlouroMyelin 6-9, GFP/dMBP 6-12. Sections counts were not averaged across individual animals to focus on comparisons on an individual section/region level which allowed for analysis of enrichment within lesions compared to non-lesion white matter. Lesion to non-lesion cell density ratios were generated by calculating the ratio of average lesional density/average non-lesional density for each spinal cord transverse section. For remyelination reporter assessment *Pdgfra-Cre^ERT^*;*Tau^mGFP^*^+/-^ ventral medial lumbar spinal cord stained with GFP/MBP/NF were imaged on W1 SoRA spinning disk confocal with Nikon NIS-Elements using 40x oil objective (108nm/pixel) with z-stacks and maximum intensity projections generated for ROIs.

### Transmission electron microscopy (TEM)

Young (P131, 4-month-old) and aged (P366-P426, 12-14-month-old) C57BL/6J (Jackson Laboratory 000664) wild-type mice and *Pdgfra-Cre^ERT^*hemizygote +/- (Jackson Laboratory 018280); *Tau^mGFP^* hemizygote +/- (Jackson Laboratory 021162) were used for transmission electron microscopy experiments. C57BL/6J mice were perfused with 4% PFA/5% glutaraldehyde and *Pdgfra-Cre^ERT^*;*Tau^mGFP^*mice for immunogold were perfused with 4% PFA/0.25% glutaraldehyde. On day 10 post-adoptive transfer young and aged EAE mice were anesthetized with intraperitoneal injection of ketamine (90mg/kg)/ xylazine (10mg/kg) and perfused transcardially with 15ml of 0.1M PBS followed by cold fixation solution: PFA (EMS 15714-S) and glutaraldehyde (EMS 16210) in 80mM sodium cacodylate and 9mM calcium chloride dihydrate in EM grade dH_2_O (EMS 22800-01) pH 7.4. The spinal cord was dissected out of the column and post-fixed for 24 hours in perfusion buffer (2 hours for immunogold followed by switch to 1% PFA overnight). Spinal cord was cut into 2mm transverse segments on spinal cord matrix (EMS 69085-C) and 3-5 lumbar and 3-5 cervical blocks per animal were washed with wash buffer (70mM sodium cacodylate, 70mM sodium chloride, 0.5mM calcium chloride, 1mM magnesium chloride hexahydrate in EM grade dH_2_O) for 30 minutes followed by post-fixation in 1% osmium tetroxide (EMS 19152) in 100mM sodium cacodylate for 1 hour protected from light. Blocks were washed with EM grade dH_2_O twice 2 minutes each then transferred to glass vials followed by dehydration (50% EtOH, 70% EtOH, 80% EtOH, 95% EtOH, 100% EtOH x 2, 100% propylene oxide 2 times, with each wash 30 minutes). Blocks were incubated overnight in 1:1 Embed 812 resin (EMS 14120, medium hardness concentration):propylene oxide (EMS 20412) at room temperature. Solution was changed to 3:1 Embed 812 resin:propylene oxide for 24 hours followed by 100% Embed 812 resin for 24 hours at room temperature. Embedding molds were filled with fresh Embed 812 resin and individual spinal cord blocks were transferred to molds and baked at 60°C for 24 hours. Semi-thin sections from lumbar spinal cord blocks were cut on Leica Ultracut UCT at 1μm thickness with Histo Diamond Knife (EMS 60-HIS) and stained with 1% toluidine blue to visualize lesions. Ultrathin sections for TEM were cut on Ultra Diamond Knife (EMS 40-UL) at 70nm thickness on Ultramicrotome Reichert Ultracut E and mounted onto grids (Pelco 200 mesh 1GC200). Grids were stained using Hiraoka staining kit (EMS 71560-00) with uranyl acetate alternative (EMS 22405) for 20 minutes followed by rinses in EMS grade dH_2_O 30 seconds 3 times, then incubation in 3mg/ml lead citrate in 0.1N NaOH in dH_2_O for 3 minutes. Grids were washed with 20mM NaOH followed by 2 washes with dH_2_O for 30 seconds then air dried. For immunogold grids were stained with Hiraoka staining kit washed with 0.01% Triton TBS, blocked with 1% NGS (JacksonImmuno 005-000-121), 1% BSA (Sigma A4161), 1% fish gelatin (Sigma G7041) in TBST (0.01% Triton TBS) for 30 minutes then incubated with anti-GFP (Abcam ab13970, 1:100) overnight at 4°C. Grids were washed with TBST 3 times, incubated in 12nm colloidal gold donkey anti-chicken (JacksonImmuno 703-205-155) in blocking solution for 90 minutes at room temperature. Grids were washed with TBST, post-fixed in 1% glutaraldehyde in PBS for 5 minutes, washed in EM grade dH_2_O followed by staining for myelin as described above. Grids were imaged on FEI Tecnai G2 Spirit TEM and ventral side was identified by the presence of ventral central vein and ventromedial white matter regions were imaged at 1150x magnification. All lumbar blocks were sectioned, imaged and quantified. All lumbar blocks from aged and young animals contained lesions with notable loss of myelin and presence of immune infiltrate in this region. TEM lesion images acquired at 1150x resolution representing one grid box were analyzed in MyelTracer (54) with users selecting and manually outlining myelin sheaths for all axons with ≥2μm given the difficulty in outlining smaller diameter axons at this magnification and desire to quantify remyelination which is more feasible for large diameter axons. Axon diameter and axon plus myelin diameters and calculated g-ratios were exported and compiled in excel and g-ratios were plotted with a cutoff of 2μm axon diameter in GraphPad Prism. For quantification of thin myelin sheaths with or without degenerative axon components TEM lesion images acquired at 1150x resolution were inverted in ImageJ/Fiji for highlighting myelin sheaths and degenerative axon components. Axons with g-ratio of 0.8 or less with a high number of mitochondria, myelin infoldings, unraveling or blebbing, and/or electron dense material were counted as thinly myelinated axons with degenerative features. Axons with g-ratio of 0.8 or less without these features were counted as without degenerative features and likely remyelinated axons. The percentage of these two types of axons was quantified over total myelinated axons with 2μm or greater axon diameter for each 1150x lesion TEM image. Total myelinated axons with 2μm or greater axon diameter were counted by using StarDist ImageJ/Fiji plugin followed by manual confirmation and adjustment as needed.

### Flow cytometry

Transgenic *Pdgfra-Cre^ERT^*; *RCE- Rosa26^flstopflEGFP^*mice that received tamoxifen and Th17 T cells or no Th17 T cells (naïve) as described above were sacrificed at 9 days post adoptive transfer for flow cytometry analysis of GFP reporter expression. Mice were anesthetized with intraperitoneal injection of ketamine (90mg/kg)/xylazine (10mg/kg) and perfused transcardially with 15ml of 0.1M PBS. Brains and spinal cord were dissected and dissociated in 20U/ml papain and 100U/ml of DNase in papain buffer (1mM EDTA, 5mM L-cystine, EBSS pH 7.4-7.6) for 30-45 minutes in 37°C 5% CO_2_ incubator with 3-4 manual trituration steps during incubation to dissociate into single cell suspension. Dissociated cells were spun at 800g 5min 4°C and resuspended in 30% Percoll (VWR 89428-524) in FACs buffer (2% FBS in PBS no cations) and spin 1000g 10 minutes 4°C speed 8. Myelin layer was removed and pellet was resuspended in FACs buffer followed by transfer over 70mm filter, PBS wash and resuspension in PBS with Fc block (BD Biosciences rat anti-mouse CD16/CD32 552142, 1:200) and viability stain (eBioscience 65-0865-14, 1:1000) for 10 minutes on ice. Cells were washed in FACs buffer and resuspended in primary antibody CD45 e450 (Invitrogen 48-0451-82 1:100) in FACs and incubated 30 minutes on ice followed by wash and resuspension in FACs. Flow cytometry of spinal cord and brain samples was performed with BD FACS Melody 4-way sorter and analyzed in FlowJo software.

### Statistics and reproducibility

GraphPad Prism was used to generate graphs and perform statistical analysis. EAE scoring and quantification was blinded by use of genotyping animal ID that was later linked to a sample group in GraphPad prism analysis. A target sample size of 6 mice per group for immunohistochemistry and 4 mice per group for TEM were set for our analysis. For IHC analysis, average EAE score at sacrifice for young was 3.5 (minimum 2.5, maximum 4.5) and aged 3.8 (minimum 2.5, maximum 4.5) with male and females in both groups. A minimum of 4 animals per group and 3 sections per animal were analyzed for IHC outcomes and individual section and lesion data-points were represented to allow for more robust statistics given potential variability on a micro and macroenvironment level within different lesions and sections. For transmission electron microscopy on wild-type mice a total of 4 young and 6 aged wild-type Th17 AT mice were sacrificed and processed with an average EAE score at sacrifice for young was 2.3 (minimum 2, maximum 2.5) and aged 3.6 (minimum 2.5, maximum 4) with males and females in both groups. All lumbar spinal cord blocks were quantified from these animals. In each figure and legend, the number of biological replicates and regions of interest (ROI) quantified, statistical test and significance levels are indicated. Due to complexity of variability including recombination, EAE severity, region, lesional difference and age; datasets did not meet normality assumptions. Statistical significance was assessed using nonparametric Mann-Whitney (two-tailed) and 2-sample median bootstrap (100,000 resamples, two-tailed, shift bootstrap, BCa bias-corrected & accelerated), with bootstrap preferred with better control for Type I errors and unequal variances (61).

## Study approval

All animal experiments were conducted in accordance with the National Institute of Health Guide for the Care and Use of Laboratory Animals and were approved by The Ohio State University Institutional Animal Care and Use Committee (IACUC) protocol 2023A00000086.

## Data availability

Data presented in available in the Supporting Data Values file. All macro codes have been made publicly available in GitHub-https://github.com/coleayden/spinalcordmacros. For additional data inquires contact the corresponding author

## Author contributions

CAH conceptualized the project. WS, MAW, and BB provided scientific guidance. EEF and CAH conducted the experiments. EEF, CNB, NFA and BJT performed quantification of histology. BJB, EEF, CNB, and NFA performed quantification of transmission electron microscopy. DP developed initial macro code and CAH modified macro codes. CAH wrote the manuscript. All authors contributed to editing and finalizing the manuscript.

## Acknowledgements

We would like to thank the following contributors: Dr. Benjamin M. Segal for scientific mentorship and access to shared equipment used for this study. Dr. Dana McTigue and Dr. Anthony Brown for assistance with transmission electron microscopy protocols and equipment usage. For equipment used for TEM sectioning and TEM imaging we acknowledge resources from the Campus Microscopy and Imaging Facility (CMIF) and the OSU Comprehensive Cancer Center (OSUCCC) Microscopy Shared Resource (MSR), The Ohio State University. This facility is supported in part by grant P30 CA016058, National Cancer Institute, Bethesda, MD. We would like to thank Jeffery Tonniges CMIF director for providing personal training for TEM ultrathin sectioning.

