## Supplemental Figure 1 for "Oligodendrocyte progenitor cell responses to inflammatory demyelination with aging in mouse model of multiple sclerosis"

A

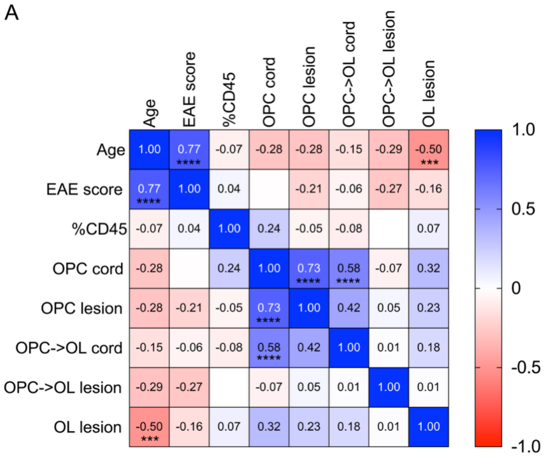


Supplemental Figure 1- Correlation analysis of OPC/OL densities, age, EAE score and CD45 infiltrate across spinal cord sections

**(A).** Spearman correlation analysis of OPC/OL manual counts on an individual spinal cord section level for variables including OPC (GFP+Aspa-CD45-), OPC->OL (GFP+Aspa+CD45-), OL (GFP-Aspa+CD45-) within the whole cord and mean of pooled lesions for cord cross-section as well as degree of CD45 infiltrate within the section, EAE score and age. Positive and negative correlations with Spearman $r<-0.5$ or $>+0.5$ and significant p value of $\leq 0.001$ are indicated with notation of p value in box. ***$p \leq 0.001$, ****$p<0.0001$. n= 48-55 spinal cord transverse sections.
